# Quantifying the nuclear localisation of fluorescently tagged proteins

**DOI:** 10.1101/2024.10.31.621290

**Authors:** Julien Hurbain, Pieter Rein ten Wolde, Peter S. Swain

## Abstract

Cells are highly dynamic, continually responding to intra- and extracellular signals. Monitoring and measuring the response to these signals at the single-cell level is challenging. Signal transduction is fast, but reporters for downstream gene expression are typically slow, requiring fluorescent proteins to be expressed and to mature. An alternative is to fluorescently tag and then monitor the intracellular locations of transcription factors and other effectors. These proteins move in or out of the nucleus in minutes, after upstream signalling modifies their state of phosphorylation. Although such approaches are being used increasingly, there is no consensus on how to quantify the nuclear and cytoplasmic localisation of these proteins. Using budding yeast, we developed a convolutional neural network that quantifies nuclear localisation from fluorescence and, optionally, bright-field images. Focusing on the cellular response to changing glucose, we generated ground-truth data using strains with both a transcription factor and a nuclear protein tagged with fluorescent markers. We then showed that the neural network based approach outperformed seven published methods, particularly when predicting single-cell time series, which are key to determining how cells respond and adapt. Collectively, our results are conclusive — using machine learning to automatically determine the appropriate image processing consistently outperforms ad hoc methods. Adopting such methods promises to both improve the accuracy and, with transfer learning, the consistency of single-cell analyses.

## Introduction

Cells continually have to respond and adapt to changes in their environment. These changes are typically detected and relayed via signal transduction networks, which activate or deactivate transcription factors, leading to altered gene expression (Fig. 1A). In eukaryotes, changes in their activation state can cause the transcription factors to either enter or exit the nucleus (Vandromme *et al*., 1996). Examples in mammalian cells include NF-*κ*B (Hoffmann *et al*., 2002; Tay *et al*., 2010) and p53 (Lahav *et al*., 2004) and at least tens of transcription factors translocate in budding yeast (Conrad *et al*., 2014).

**Fig. 1.**
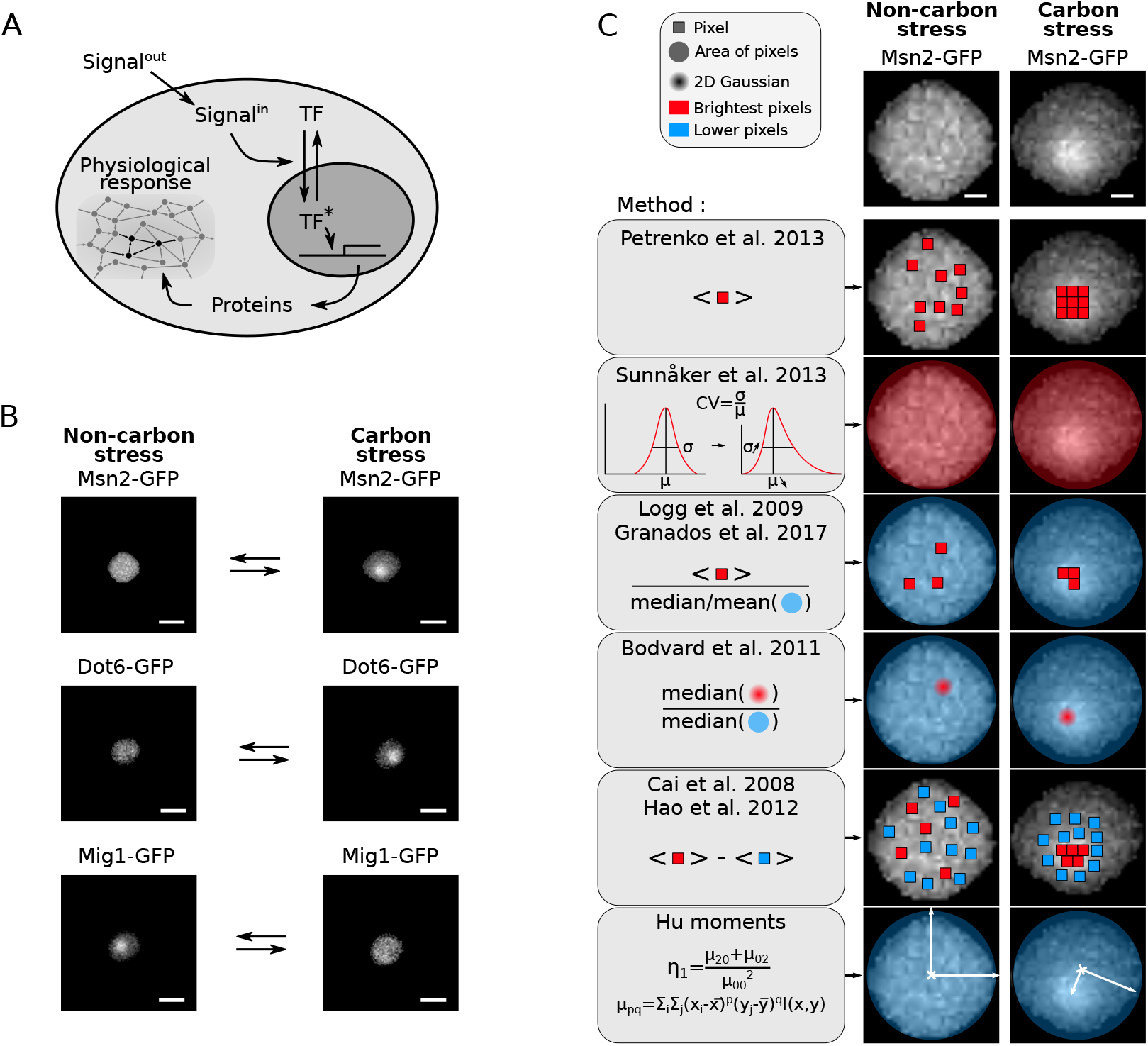
Detecting and quantifying the nuclear translocation of transcription factors. (A) A schematic showing a transcription factor entering the nucleus, regulating gene expression, and so cellular physiology through the proteins synthesised. *Signal*^*out*^ and *Signal*^*in*^ are the extracellular and intracellular signals that generate active transcription factor. (B) The responses of three transcription factors in budding yeast to changes in extracellular glucose. Msn2 is the master regulator of the general stress response; Dot6 is a repressor of ribosome biogenesis; and Mig1 represses regulons for metabolising alternative sugars to glucose. Scale bar = 4*µm*. (C) Seven methods to quantify the localisation of transcription factors. Left and right images show Msn2-GFP in a single cell. We indicate the pixels used by each method with colours, those with higher fluorescence in red and those with lower in blue. Areas and 2D Gaussian use ensembles of pixels. The first Hu moment, *η*_1_, is a function of the pixels intensities at positions *x* and *y*. Scale bar = 1*µm*. (B-C) Images in non-carbon stress are for 2% glucose; those in carbon stress are for 0% glucose.

Monitoring these transcription factors as they translocate is a versatile method to characterise how cells respond to changes in their environment or intracellular state. It can be applied to single cells, to time-varying environments and to dynamic cellular responses, and has a time resolution of minutes, much closer to that of the underlying signal transduction than reporters of gene expression.

For budding yeast, following the nuclear translocation of transcription factors and other proteins involved in regulating transcription has become a standard way to visualise and characterise stress responses over time. For example, the high-osmolarity glycerol (HOG) MAP kinase is often monitored to investigate hyper-osmotic stress (Hersen *et al*., 2008; Mettetal *et al*., 2008). Whereas a fall in extracellular glucose generates nuclear entry of both Msn2 (Jacquet *et al*., 2003; Hao *et al*., 2013), a master regulator of the general stress response (Gasch *et al*., 2000; Causton *et al*., 2001), and of Dot6 (Gasch *et al*., 2017; Granados *et al*., 2018), a repressor of ribosome biogenesis (Lippman and Broach, 2009). Simultaneously Mig1 (Bendrioua *et al*., 2014; Lin *et al*., 2015), a glucose-sensing specialist and a repressor of regulons for metabolising other sugars (Conrad *et al*., 2014), exits the nucleus (Fig. 1B).

Although all the approaches developed use fluorescent-protein tags to identify transcription factors, they differ in how they quantify their intracellular localisation (Fig. 1C). To avoid generating strains with a second fluorophore marking the nucleus, measuring the spatial homogeneity of the transcription factor’s fluorescence signal is often used as a proxy for nuclear localisation — a cytoplasmic transcription factor generates a fluorescence spread more equally over the cell than one concentrated in the nucleus (Fig. 1B). Nevertheless, there is no consensus on how to measure spatial inhomogeneity, with at least seven different methods proposed for transcription factors in yeast alone (Table 1).

**Table 1.** Methods to quantify the nuclear localisation of transcription factors from single-cell fluorescence images.

| Name | Method | Reference |
| --- | --- | --- |
| Petrenko | Mean of the pixels with the 20% highest intensities | Petrenko *et al.* 2013 (Petrenko *et al.*, 2013) |
| Sunnåker | Coefficient of variation of the pixel intensities | Sunnåker *et al.* 2013 (Sunnåker *et al.*, 2013) |
| Logg | Ratio of the mean of the three highest pixels to the mean of the remaining pixels | Logg *et al.* 2009 (Logg *et al.*, 2009) |
| Granados | Ratio of the mean of the five highest pixels intensities to the median of the remaining pixels | Granados *et al.* 2018 (Granados *et al.*, 2018) |
| Bodvard | Ratio of the median value of a smoothed Gaussian centred on the brightest pixel over the median of the remaining pixels | Bodvard *et al.* 2011 (Bodvard *et al.*, 2011) |
| Cai Hao | Difference of the mean of the five brightest pixels and the mean of the remaining pixels | Cai *et al.* 2008 (Cai *et al.*, 2008)<br>Hao *et al.* 2012 (Hao and O'Shea, 2012) |
| Hu moments | First invariant moment $\eta_1$ | Hu, M.-K. 1962 (Hu, 1962)<br>Nasrudin *et al.* 2021 (Nasrudin *et al.*, 2021) |
| Neural network | Prediction by a regression convolutional neural network | This study |

We set out to evaluate the different methods, comparing these approaches with one that we developed using a convolutional neural network (Goodfellow *et al*., 2016), as well as with an older standard for measuring spatial inhomogeneities: Hu moments (Hu, 1962; Nasrudin *et al*., 2021). These moments use the moments of inertia treating the fluorescence values within a cell as masses.

## Results

### Finding transcription-factor localisation from a nuclear marker

We generated time-lapse single-cell data with both a fluorescent transcription factor, tagged with GFP, and a fluorescent nuclear marker, the Nhp6a gene tagged with mCherry (Hansen *et al*., 2015; Granados *et al*., 2018) (Fig. 2A). The Nhp6a protein remodels nucleosomes (Ruone *et al*., 2003). We used the ALCATRAS microfluidic device to grow and trap cells (Crane *et al*., 2014) and the BABY algorithm (Pietsch *et al*., 2023) to segment and track these cells over time from time-lapse, bright-field images.

**Fig. 2.**
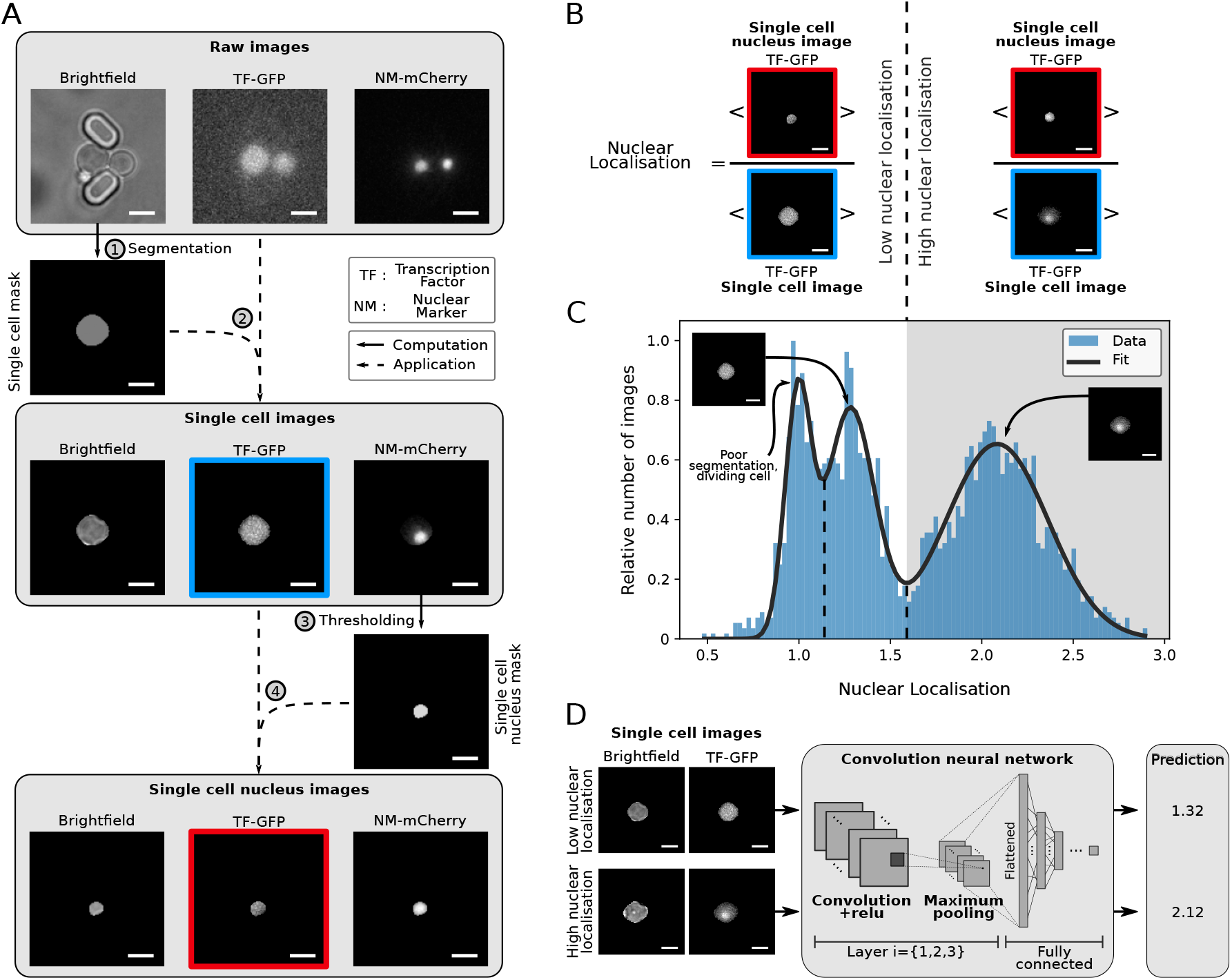
Estimating the nuclear localisation and preparing images for analysis by a neural network. Scale bar = 4*µm*. (A) Our pipeline for preparing images for analysis. Cells grow stably between two pillars in the ALCATRAS microfluidic device (Crane *et al*., 2014), which has hundreds of these traps. Using the BABY algorithm (Pietsch *et al*., 2023), we segment single cells from the bright-field images and find the pixels corresponding to these cells in the fluorescence images. With Otsu thresholding of the image of the nuclear protein Nhp6a-mCherry, we determine the pixels in the nucleus. For each cell, we therefore have a bright-field image, a fluorescence image showing the intracellular fluorescence from a tagged transcription factor, and a binary image identifying the nuclear pixels. (B) A schematic showing the definition of nuclear localisation, Eq. 1. (C) The distribution of nuclear localisation typically has three peaks. We show the distribution for Msn2-GFP combining images from a time-lapse experiment where we switched extracellular glucose from 2% to 0%. The images are from just before and after the switch. (D) The convolutional neural network has three layers and predicts nuclear localisation using the maximum projection over Z stacks of both fluorescence and bright-field images as inputs.

To identify the nucleus from the mCherry images, we used Otsu thresholding (Otsu, 1979). Just as there is no consensus on the optimal way to measure the localisation of transcription factors, there is none too on how to determine the nucleus’s location (Jacquet *et al*., 2003; Duveau *et al*., 2024; Li *et al*., 2018). In contrast to transcription factors, however, the fluorescence from a nuclear marker is consistently bright over both time and extracellular conditions (Fig. 1C), and we found little difference between most methods (Fig. S1), all of which threshold the image in some way (Kapur *et al*., 1985; Johannsen and Bille, 1982; Bernsen, 1986; Parker, 2010). We chose Otsu thresholding because it has no parameters; it does though require a bimodal distribution of pixel fluorescence intensities (Goh *et al*., 2018) and so a sufficiently bright nuclear marker. These conditions were satisfied in our experiments.

With the nucleus identified, we defined, following Duveau *et al*. (2024), the degree of nuclear localisation as

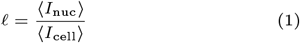

where *I*_nuc_ represents the pixel intensities in the nucleus and *I*_cell_ those for the whole cell, including the nucleus (Fig. 2B). The most relevant quantity for predicting gene expression is however the nuclear concentration ⟨*I*_nuc_⟩, the total fluorescence in the nucleus divided by its area. We can recover this concentration from the localisation by multiplying by ⟨*I*_cell_⟩, the mean cellular fluorescence, which is commonly measured.

Applying our workflow to cells in different carbon sources, we typically found a tri-modal distribution of the level of nuclear localisation (Fig. 2C). The largest mode corresponds to cells with a high nuclear concentration of transcription factor; the middle node to cells with a low nuclear concentration; and the lowest mode, based on visual inspection of the images, either to poorly segmented cells, such as those overlapping the edges of an image, or to cells actively dividing with consequently an imperfectly identified nucleus. We therefore consider cells with predominantly cytoplasmic transcription factors to have 1.15 *<* 𝓁 *<* 1.65 and cells with predominantly nuclear transcription factors to have 𝓁 *>* 1.65.

### A convolutional neural network to predict nuclear localisation

We developed a convolutional neural network (Goodfellow *et al*., 2016) to predict nuclear localisation from segmented fluorescence images of tagged transcription factors (Fig. 2D). The network comprises three layers and a fully connected layer (Fig. S2A) and maps microscopy images of a single cell to one continuous value, the predicted level of nuclear localisation. As inputs, we used the maximum projection of fluorescence images taken at five Z sections and, although not required, the maximum projection of the corresponding five bright-field images. Such bright-field images are readily available and improved accuracy because the nucleus can sometimes be partly seen in bright-field (Adjavon, 2022). We used the nuclear Nhp6a-mCherry marker only to establish a ground-truth data for training; the corresponding images were not used as network inputs. After training, using data from the transcription factors Msn2, Dot6, and Mig1, the network had an accuracy of 95% (Fig. S2B). Excluding data from one specific transcription factor while training but not while testing neither substantially nor consistently affected performance (Fig. S2C). For instance, removing data from Msn2 while training does not always lead to poor performance while testing with that data.

### Comparing different methods

With the nuclear marker determining the ground-truth level of nuclear localisation, we compared the various methods using three transcription factors that respond to extracellular glucose (Fig. 3A). Two of these enter the nucleus in low glucose while the other exits (Fig. 1B). A difficulty is that each method makes predictions over a different numerical range because each has its own way of characterising spatial inhomogeneity. We therefore plotted the log_2_ ratio of the method’s predicted value to the ground-truth level of nuclear localisation and centred the results by subtracting each method’s mean ratio (non-centred results are in Fig. S3). Accurate methods should have a tight, symmetric distribution.

**Fig. 3.**
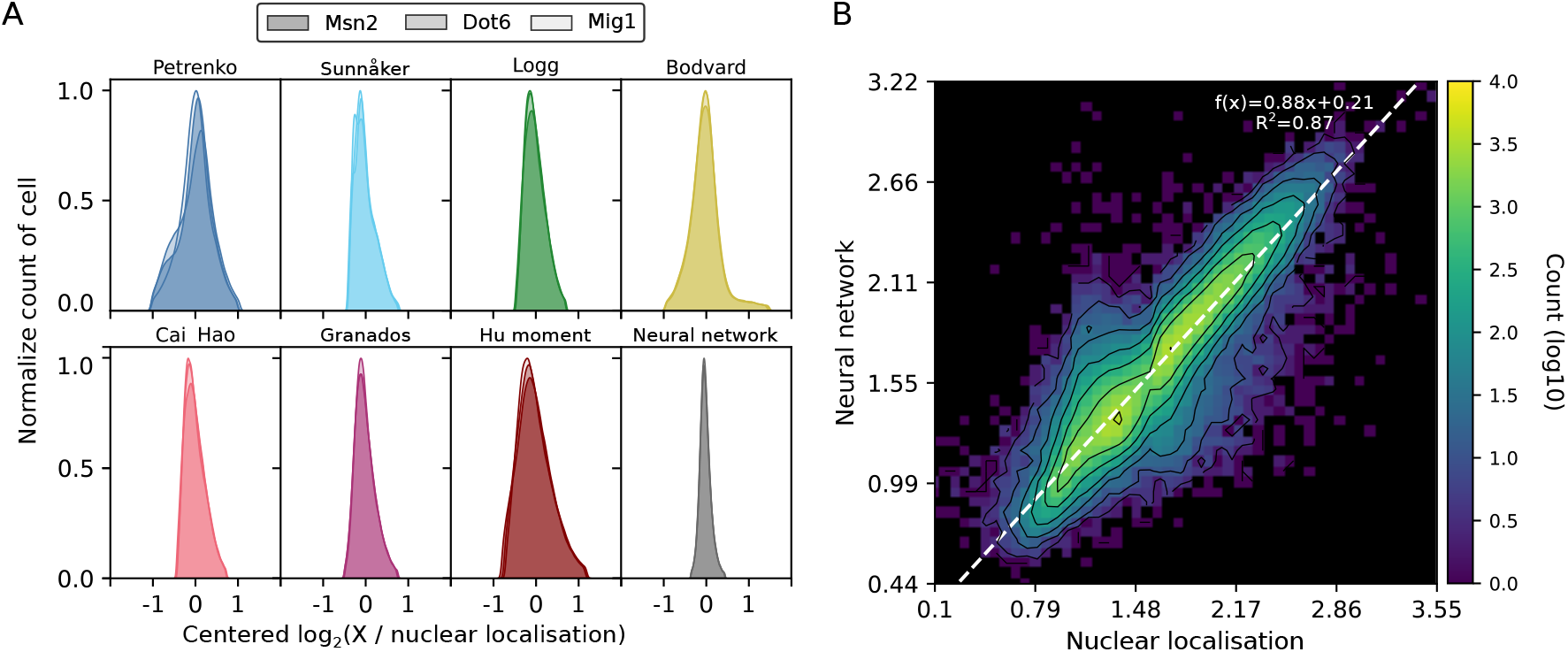
The neural-network approach best identifies nuclear localisation from among the different methods. (A) The neural network has the tightest distribution of errors when comparing predictions to the ground-truth level of nuclear localisation found from the Nhp6A-mCherry marker. Data are for Msn2-GFP, Dot6-GFP, and Mig1-GFP from cells in a step from 1% to 0% extracellular glucose. We plot the log_2_ of the predicted to the ground-truth value and subtracted the mean over all cells, so that each method has an identical mean of zero. For the approach of Bodvard *et al*. (Bodvard *et al*., 2011), we ensured all values are positive by incrementing each by one. (B) Plotting the neural network’s prediction against the ground-truth level of nuclear localisation shows a tight linear relationship. The shading indicates numbers of cells on a *log*_10_ scale, with zero cells in black. The corresponding plots for the other methods are in Fig. S4.

The neural network performed best. Although having a skewness comparable to other approaches (Fig. S3), it had the lowest standard deviation and highest consistency over the three transcription factors. Plotting the predicted localisation versus the ground truth (Fig. 3B), the neural network’s results tightly followed the *y* = *x* line (Fig. S4), and both its correlation and mutual information with the ground truth were highest (Fig. S5). Unlike the other methods, as it is the only one trained to do so, we can also directly interpret its predictions as estimates of Eq. 1.

Although the neural network’s performance is perhaps unsurprising, being the only method benefiting from the training data, it is useful to quantify how much better it is over the alternatives. To this end, we investigated time-series — the focus of multiple studies (Bendrioua *et al*., 2014; Cai *et al*., 2008; Duveau et al., 2024; Goulev et al., 2017; Granados et al., 2017, 2018; Hansen and O’shea, 2013; Hao et al., 2013; Hersen *et al*., 2008; Lin *et al*., 2015; Mettetal *et al*., 2008; Pietsch *et al*., 2023). The results further highlight how improved the predictions can be.

### The neural network’s predictions best capture dynamic single-cell responses

The variation in how individual cells respond and its information often manifests neither in the response at a given moment in time nor in the long-time limit, but in the dynamical trajectory of the response over time (Tostevin and Ten Wolde, 2009; Granados *et al*., 2018). In constant glucose (Fig. 4), both Msn2 and Dot6 transiently enter the nucleus (Dalal *et al*., 2014), impeding straightforward averaging, which would obscure such behaviour. Almost all cells responded however when we removed extracellular glucose, with Msn2 entering the nucleus on average faster than Dot6.

**Fig. 4.**
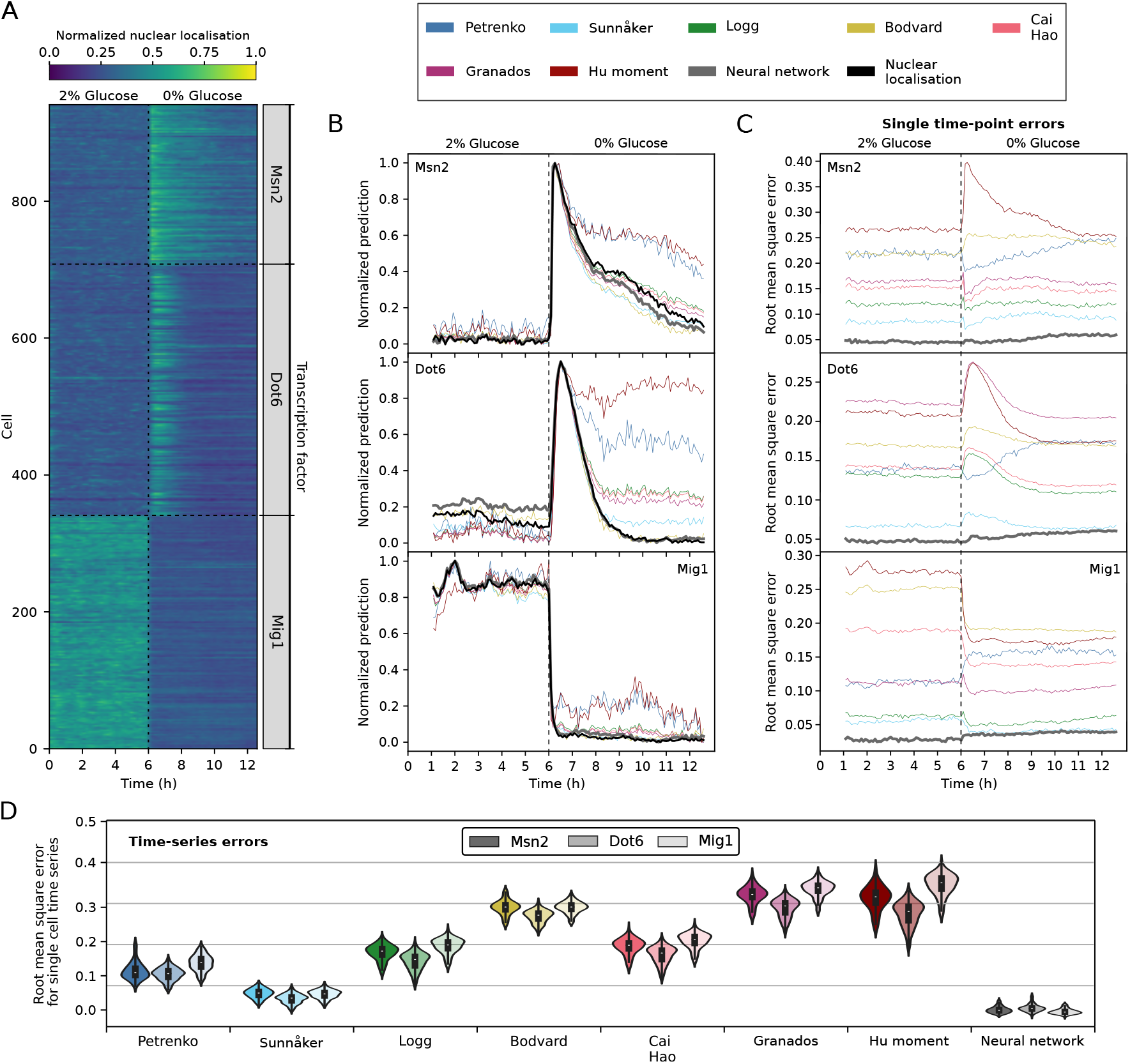
The neural network best predicts single-cell time-series of nuclear localisation. (A) The ground-truth time series of nuclear localisation for cells experiencing a drop in glucose from 1% to 0% after six hours of growth. Each row represents a single cell with the shading showing the normalised degree of nuclear localisation. For Msn2, *n* = 233; for Dot6, *n* = 360; for Mig1, *n* = 341. (B) The predicted mean level of nuclear localisation over all cells at each time point varies between the different methods (predictions for individual cells are in Fig. S6). The grey curve corresponding to the neural network is so close to the ground truth that it is often obscured. We normalised each prediction to be between zero and one. For Petrenko *et al*.’s method (Petrenko *et al*., 2013), we inverted their prediction so that it behaves similarly to the others. (C-D) The root mean square error is defined as the root mean square of the difference between the ground-truth nuclear localisation and that identified by the method. (C) The root mean square errors at each time point for all cells. (D) The distributions of root mean square errors for each cell’s entire time series — each cell is tracked during the experiment. The root mean square error is calculated based on the predictions made at every time point where the cell is present. This process is repeated for each cell, resulting in a distribution of root mean square error values. Results for Msn2 are on the left and for Mig1 on the right.

Applying the methods to our data, we saw substantial differences even in predicting the mean nuclear translocation over time (Fig. 4B). Although all methods identified the cells as responding to the drop in extracellular glucose, shown by a peak in the time series, some had difficulty when the transcription factors were predominately in the nucleus, with higher errors for Msn2 and Dot6 after glucose drops and for Mig1 before. Only the neural network’s predictions had low errors both at each time point (Fig. 4C) and for each single-cell time series (Fig. 4D).

## Discussion

Our study shows that machine learning, through the convolutional neural network we developed, consistently outperforms *ad hoc* approaches, particularly for single-cell time series. Such time series best characterise individual responses (Murugan *et al*., 2021).

Although neural networks require training data, so too do *ad hoc* approaches because without such data these approaches cannot be validated. The difference lies in the quantity of training data, but with microfluidic technology, generating sufficient data is possible in a single experiment, as we demonstrated here. Additionally, the quality of the approach can be continually improved by adding more images to the training data, such as from different microscopes, imaging protocols, strains, or species. Even better, a network trained in one laboratory will likely need much less data to be re-trained for another, with the promise to improve the consistency of signal-cell analyses broadly.

Single-cell data is often noisy, and a poor choice of signal can further reduce the signal-to-noise ratio, confounding analyses. For nuclear translocations at least, we have shown that using machine learning to identify the signal, through optimising the image processing required, is one way to gain accuracy and with it, no doubt, greater biological insight for the future.

## Methods

### Strains

Details of all strains are given in Granados *et al*. 2018 (Granados *et al*., 2018). We grew cells in synthetic complete (SC) medium at 30°C either supplemented with either 2%, 1% glucose or with no additional carbon source, denoted 0% glucose.

#### Cell preparation and loading ALCATRAS

We used overnight cultures in a 30°*C* incubator under agitation to generate mid-log cells which we diluted in fresh SC medium to an optical density of 0.1 and then incubated for a further three hours before loading into a microfluidic device. To expose different strains to the same extracellular conditions, we used a multi-chamber version of ALCATRAS (Crane *et al*., 2014). Prior to loading, the ALCATRAS chambers were pre-filled with growth medium supplemented with 0.05% bovine serum albumin to facilitate cell loading and reduce cell clumping.

### Time-lapse microscopy

Our microscope is a Nikon Ti Eclipse, optimized for imaging GFP, mCherry, and Cy5 fluorescence, as well as bright-field transmission. For GFP, we used a Chroma dual-band filter set (59022) with an excitation range of 452-490 nm, centered at 470 nm, a beam splitter range of 496-548 nm, and an emission filter at 535/30 nm (520-550 nm range). The mCherry imaging shared the dual-band filter set for GFP, with an excitation range of 554-558 nm (centered at 556 nm), a beam splitter range of 595-677 nm, and an emission filter at 632/60 nm. Finally, for Cy5 imaging we used an excitation filter at 620/60 nm (590-650 nm, centered at 620 nm), a 660 nm longpass dichroic, and a 665 nm longpass emission filter. For Cy5 and bright-field transmission, the microscope had a white LED with a broad-spectrum range, and for fluorescence imaging, an OptoLED light source from Cairn Research.

The objective was a Nikon 60× oil-immersion lens with a numerical aperture (NA) of 1.4, using Nikon F2 immersion oil. Imaging was captured with a Teledyne Prime 95b sCMOS camera, with an 11 *µ*m pixel size, 16-bit dynamic range, 1×1 binning, gain of 1, and air cooling at −15°C. Exposure time for imaging was set at 30 ms, with images captured every five minutes for 150 time points, i.e., 12.5 hours. We acquired bright-field and fluorescence images at five Z-sections spaced by 0.6 *µ*m, but in a single focal plane for the Cy5 channel. Both the microfluidic device and the media were kept at 30°*C* inside an incubation chamber (Okolabs). Nikon’s Perfect Focus System maintained consistent focus.

We used fluigent pressure-driven system to control the flow of media. We applied carbon stress after six hours by switching the flow rate of either 2% or 1% glucose medium from 9 *µ*L/min to 1 *µ*L/min and the flow rate of 0% glucose medium from 1 *µ*L/min to 9 *µ*L/min. Slowly flowing medium did not enter the chamber containing cells and was re-directed to waste. We used Cy5 dye in the 0% glucose medium to confirm media switches. Media were supplemented with 0.05% of bovine serum albumin (BSA) to facilitate cell loading and reduce cell clumping.

### Image segmentation

To segment and track cells, we used BABY (Pietsch *et al*., 2023) and the aliby Python pipeline (Muñoz González, 2023). To identify nuclear pixels, we used Otsu thresholding on single-cell images after applying a Gaussian blur (Gonzalez, 2009). Alternative methods, which we also tested, are in Table 2.

**Table 2.** Image-processing methods to identify pixels in the nucleus from fluorescence images of a nuclear marker.

| Name | Method | Reference |
| --- | --- | --- |
| Otsu | Global thresholding | Otsu N. 1978 (Otsu, 1979) |
| Kapur | Entropy thresholding | Kapur *et al.* 1985 (Kapur *et al.*, 1985) |
| Johannsen | Entropy thresholding | Johannsen *et al.* 1982 (Johannsen and Bille, 1982) |
| Bernsen | Local thresholding | Bernsen J. 1986 (Bernsen, 1986) |
| Contrast | Local thresholding | Parker J. R. 2010 (Parker, 2010) |

We estimated mutual information with the nearest-neighbour approach (Kraskov *et al*., 2004; Ross, 2014).

### Constructing and training the neural network

The neural network has three convolutional layers (Fig. S2A). The first two comprise convolutions followed by Relu activation functions and then maximum pooling. The third layer is similar but with no maximum pooling and its output is flattened into a one-dimensional array. The final, i.e., fully-connected, layer combines a linear and Relu function.

For training (Fig. S2B), we used mean square error as the loss function and the Adam optimiser. The training data comprised ≈ 180, 000 single-cell images shuffled with a proportion of 90% for training and 10% for testing from an experiment with a switch of extracellular glucose from 2% to 0%. We used a batch size of 64 and a CUDA implementation of the network to increase speed (Garland *et al*., 2008). After some exploration, we chose a learning rate of 0.001 with a linear learning-rate scheduler of ratio 0.5 over 50 epochs. To compute accuracy, we used a nuclear localisation of 1.65 and above to define cells with a localised, nuclear transcription factor.

For all coding, we used Python and the Pytorch with CUDA package.

## Competing interests

No competing interest is declared.

## Author contributions statement

JH and PSS conceived the research; PRtW and PSS helped interpret the results and provided expertise on the methods.

JH developed the neural network, performed the simulations and experiments, and wrote the manuscript’s first draft. All authors reviewed and edited the manuscript.

## Acknowledgments

The authors thank Ivan B. N. Clark for his help in performing the experiments. This work is part of the Dutch Research Council (NWO) and was performed at the research institute AMOLF and the University of Edinburgh. This project has received funding from the European Research Council (ERC) under the European Union’s Horizon 2020 research and innovation program (Grant Agreement No. 885065). PSS gratefully acknowledges support from the BBSRC (grant number BB/W006545/1).

## Data & code availability

Codes and reduced data used in this work can be found at : https://git.ecdf.ed.ac.uk/v1jhurba/neunet-nucloc.git. The full dataset set with its corresponding code is available at : https://doi.org/10.7488/ds/7820.

## Supplementary information

**Fig. S1.**
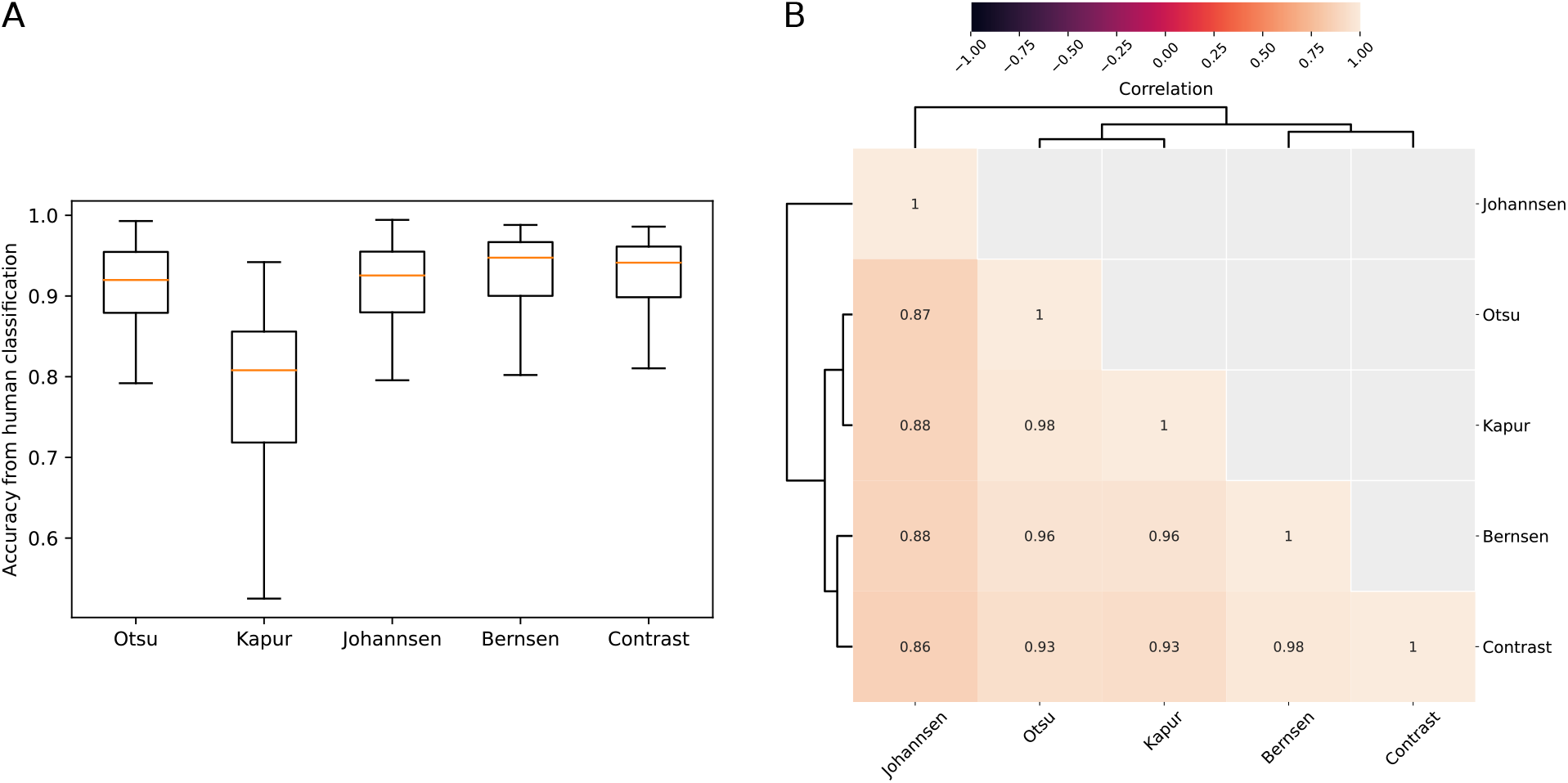
A comparison of different methods to identify the nucleus. (A) A comparison of the different methods for identifying the nucleus from a fluorescence image of a nuclear marker (Table 2). To establish a ground truth, we identified the nucleus by eye in approximately 200 images. (B) A matrix showing the high correlation between the different methods.

**Fig. S2.**
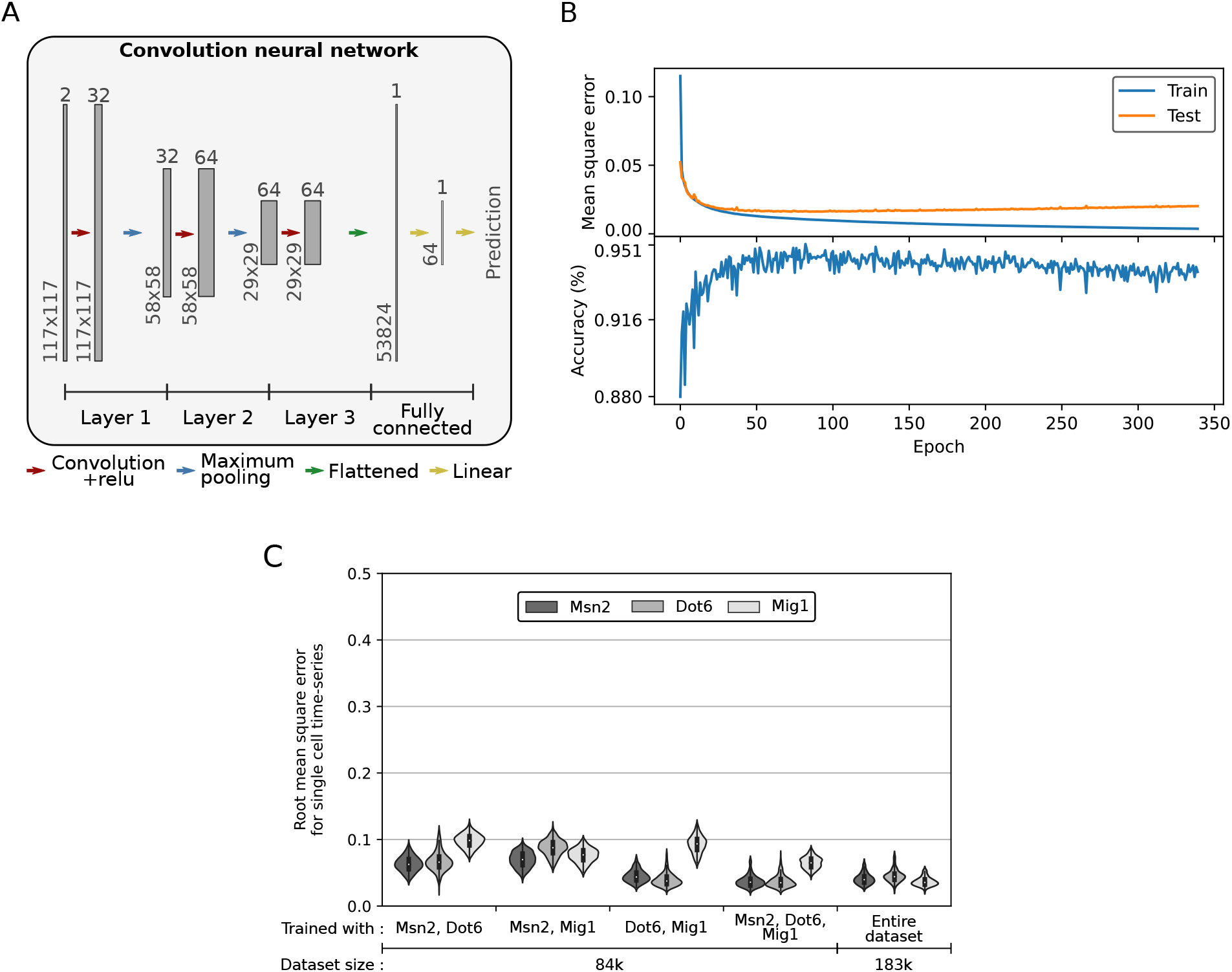
A neural network to predict localisation from fluorescence and, optionally, bright-field images. (A) The structure of the convolutional neural network. Each gray rectangle represents a different stage as the network processes the image. The size of the image is indicated on the side and the number of feature channels on top. The single-cell images we use as inputs are 117 *×* 117 pixels and have two channels: fluorescence and bright-field. (B) Typical training curves for the network. We used a mean-square loss function and ≈ 180, 000 images with 90% randomly selected for training and 10% for testing. To compute an accuracy, we applied a threshold to the predicted level of nuclear localisation (the threshold divides the white from the gray shaded areas in Fig. 2C). (C) The distributions of root mean square errors for each single-cell time series of the dataset used in Fig. 4 for networks trained on data from different pairs of transcription factors ignoring one. We chose the number of images to allow equally sized datasets for all the training conditions. The results from the entire dataset are the same as Fig. 4D. We show Msn2 are on the left and Mig1 on the right.

**Fig. S3.**
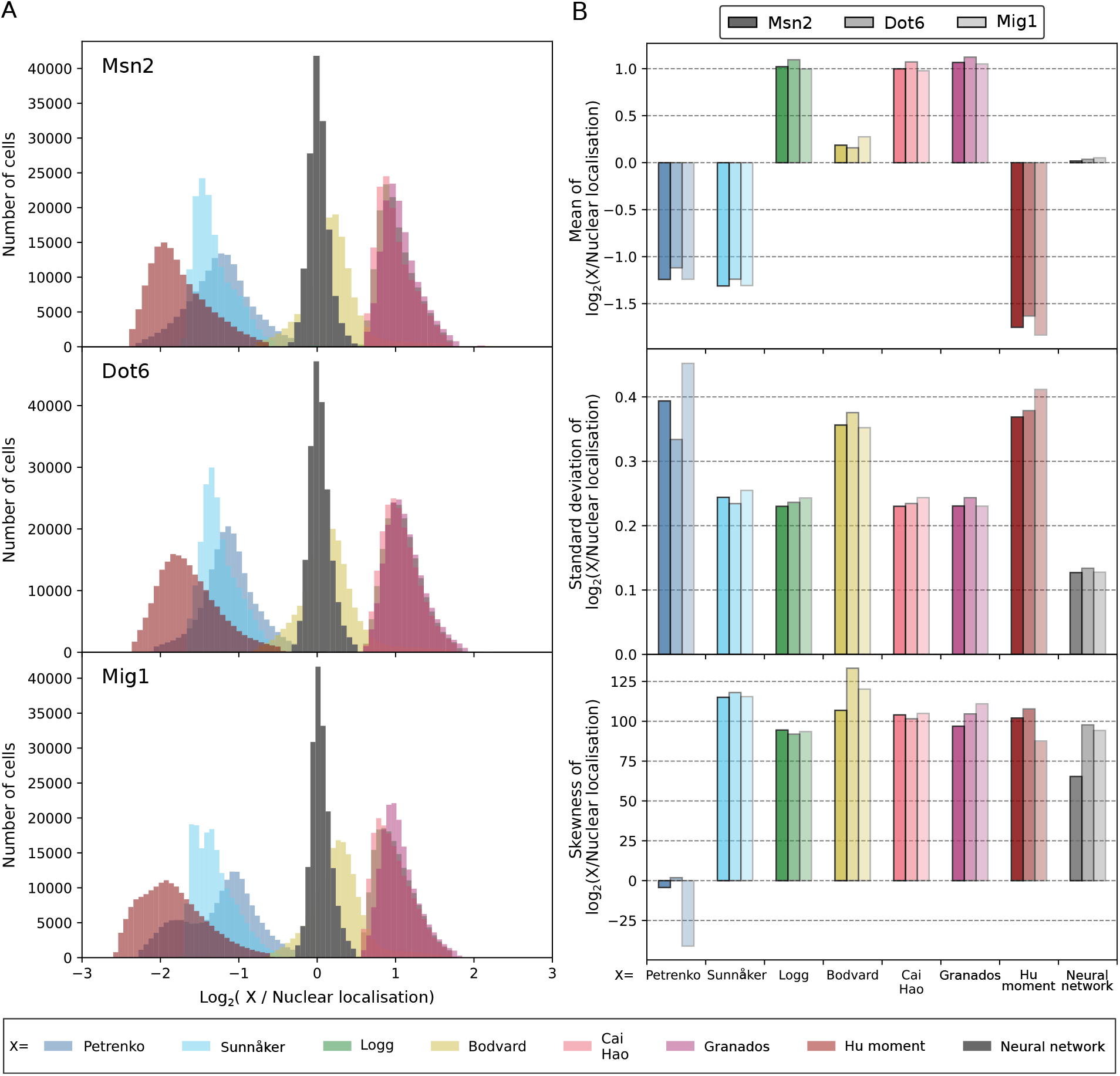
Comparing predictions of nuclear localisation on an absolute rather than relative scale. (A) The error distributions for the different methods of predicting nuclear localisation from Table 1. Unlike in Fig. 3A, these distributions are not normalised to have zero mean. Most poorly predict Eq. 1 because they are not explicitly designed to do so, although their values do correlate with the transcription factor’s localisation. We have combined data for Msn2-GFP, Dot6-GFP, and Mig1-GFP in a step from 1% to 0% extracellular glucose at all time points. For the Bodvard *et al*. method (Bodvard *et al*., 2011), we ensured values are always positive by incrementing each value by one. (B) The mean, standard deviation, and skewness of the error distributions. We use shading to indicate the different transcription factors, with Msn2-GFP on the left and Mig1-GFP on the right.

**Fig. S4.**
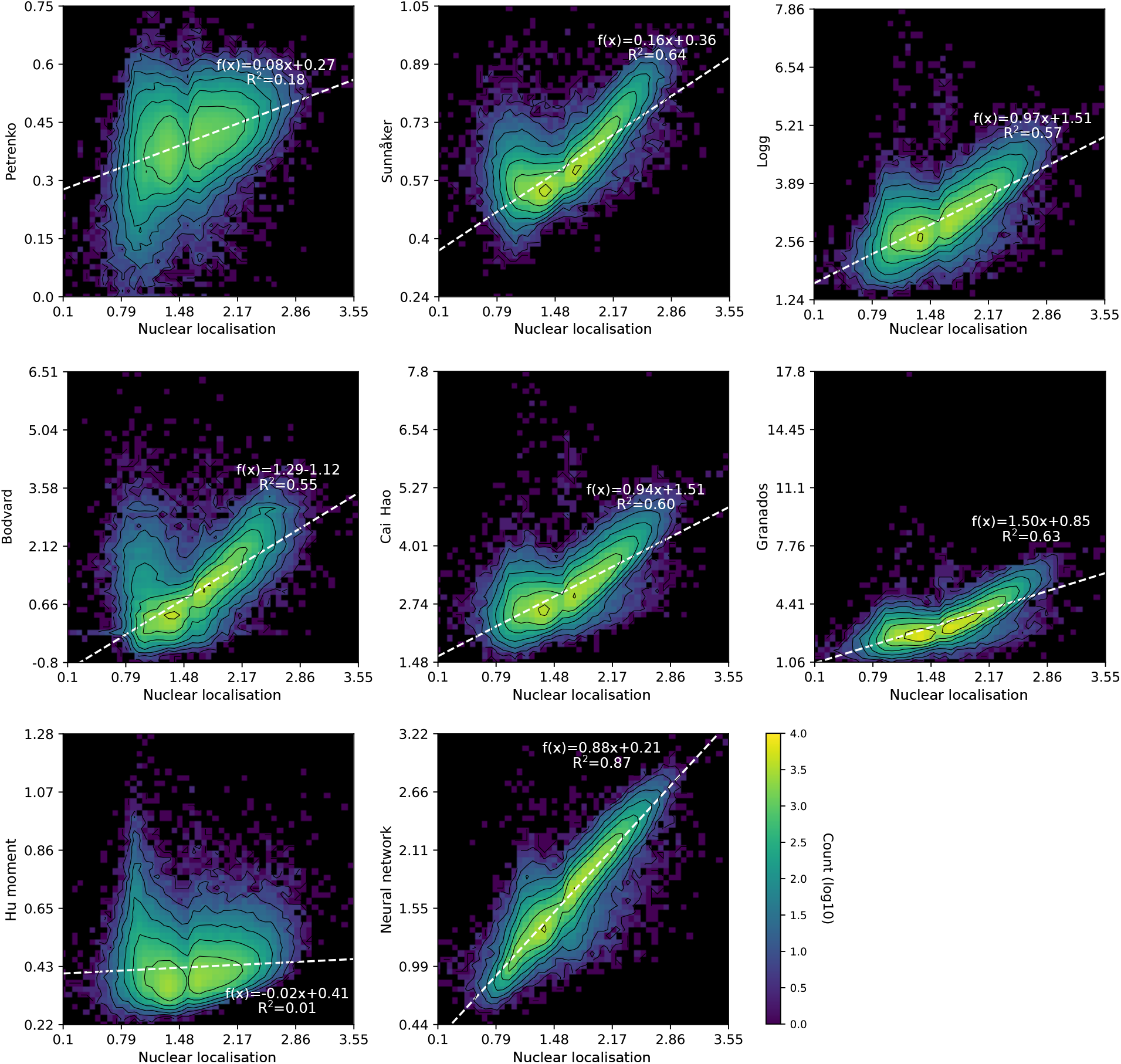
The predicted nuclear localisation versus the ground-truth localisation. Colours represent the number of cells on a log_10_ scale; black represents no cells. The white dashed-line is a linear regression. Ideal predictions should be linearly proportional to the ground truth, with a high *R*^2^.

**Fig. S5.**
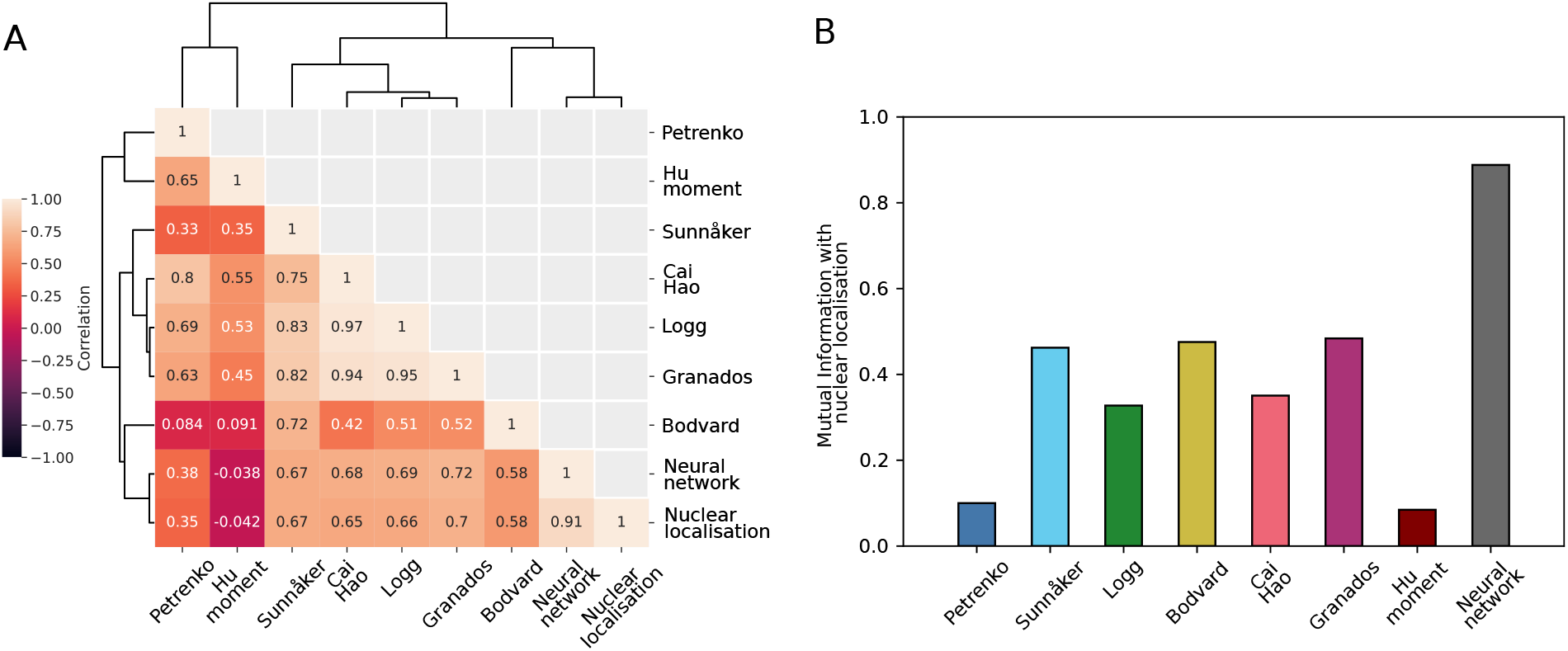
Statistically comparing the methods’ predictions to the ground-truth localisation favours the neural network. (A) A correlation matrix of the methods’ predictions. Only the magnitude of the correlation is relevant because some methods are anti-correlated with the ground truth. (B) The mutual information between the predictions and the ground-truth localisation. Unlike the correlation coefficient, the mutual information measures both linear and non-linear relationships between two variables.

**Fig. S6.**
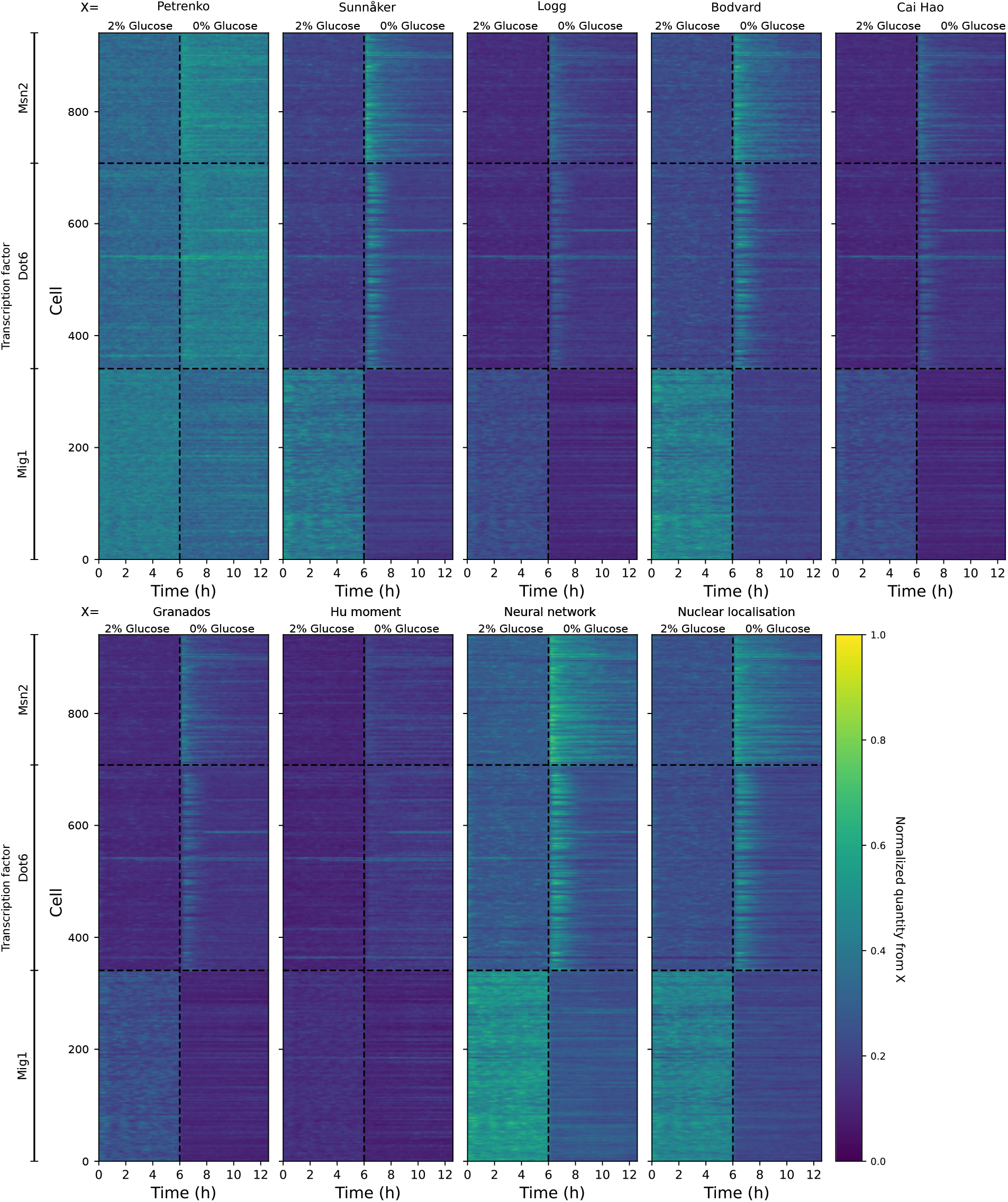
Predictions of nuclear localisation for individual cells during a drop in extracellular glucose. Each row represents a single cell, with the colour code showing the predicted level of nuclear localisation normalised to be between zero and one. We decreased the extracellular glucose concentration from 1% to 0% after six hours. To aid comparison, we inverted the values predicted by Petrenko *et al*.’s method (Petrenko *et al*., 2013), subtracting their prediction from one.

## Notes

https://git.ecdf.ed.ac.uk/v1jhurba/neunet-nucloc.git

https://doi.org/10.7488/ds/7820

### Competing Interest Statement

The authors have declared no competing interest.

